# Lack of regularity between letters impacts word recognition performance

**DOI:** 10.1101/753038

**Authors:** Sofie Beier, Jean-Baptiste Bernard

## Abstract

Physical inter-letter dissimilarity has been suggested as a solution to increase perceptual differences between letter shapes and hence a solution to improve reading performance. However, the deleterious effects of font tuning suggest that low inter-letter regularity (due to the enhancement of specific letter features to make them more differentiable) may impair word recognition performance. The aim of the present investigation was 1) to validate our hypothesis that reducing inter-letter regularity impairs reading performance, as suggested by font tuning, and 2) to test whether some forms of non-regularities could impair visual word recognition more. To do so, we designed four new fonts. For each font we induced one type of increased perceptual difference: for the first font, the letters have longer extender length; for the second font, the letters have different slants; and for the third font, the letters have different font cases. We also designed a fourth font where letters differ on all three aspects (worst regularity across letters). Word recognition performance was measured for each of the four fonts in comparison to a traditional sans serif font (best regularity across letters) through a lexical decision task. Results showed a significant decrease in word recognition performance only for the fonts with mixed-case letters, suggesting that fonts with low regularity, such as mixed-case letters, should be avoided in the definition of new “optimal” fonts. Letter recognition performance measured for the five different fonts through a trigram recognition task showed that this effect is not consistently due to poor letter identification.

## 1. Introduction

The often repeated saying among typographers that “type is a beautiful group of letters, not a group of beautiful letters” (1), suggests that it is only when letters work as a group that they become type, a visual characteristic that we name “inter-letter regularity”. To achieve this, a basic principles of sign painting and font design dictates that fonts and lettering shall be based on a repetition of shapes with the aim of ensuring harmony and balance between the letters (2, 3) (Fig 1). This means that all lower- and uppercase letters originate in two different modular systems that put together constitute the alphabet (one for lowercase letters and one for uppercase letters) (4). Such an approach naturally leads to letters of relatively similar shapes (and high regularity). By contrast, it has often been proposed that greater letter distinctiveness, where new features are added to selected letters, could facilitate reading, as it minimizes the risk of letter confusion (5-7). However, greater letter distinctiveness also decreases inter-letter regularity.

**Fig 1.**
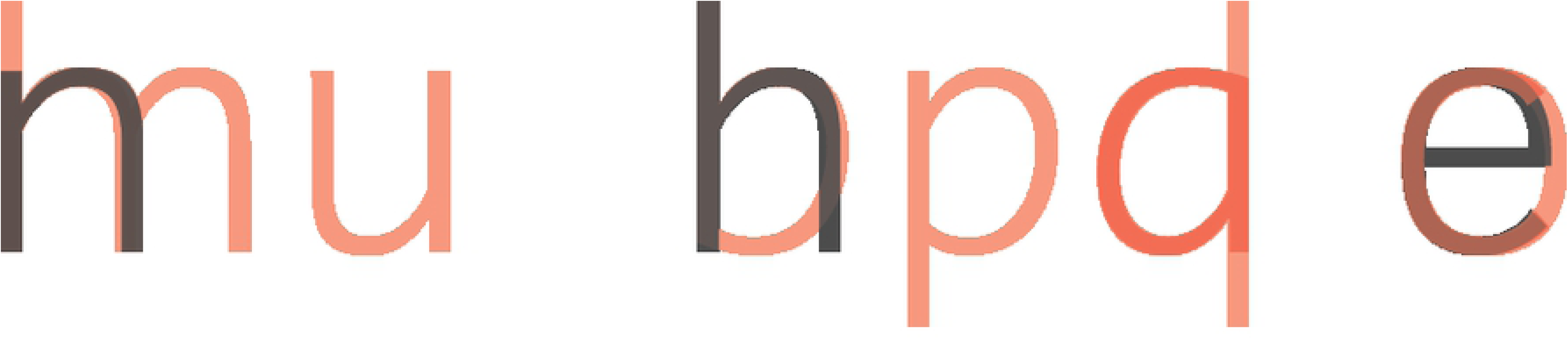
The rule of repetition of shapes in font design, here demonstrated with lowercase letters.

To investigate whether high letter differentiation could improve peripheral reading, Bernard et al. (7) created a new font, referred to as Eido (Fig 2). They found that while participants familiarized themselves with the font, their reading performance improved for both letter and word recognition, although sentence reading speed was not significantly improved. Xiong et al. (8) further found that Eido outperformed both Helvetica and Times Roman for reading acuity performance, while maximum reading speed was not significantly improved. Also interested in letter differentiation, Beier and Larson (9) measured letter recognition of variations within the same font family and found certain letter shapes of greater dissimilarity to facilitate better single letter recognition than others.

**Fig 2.**
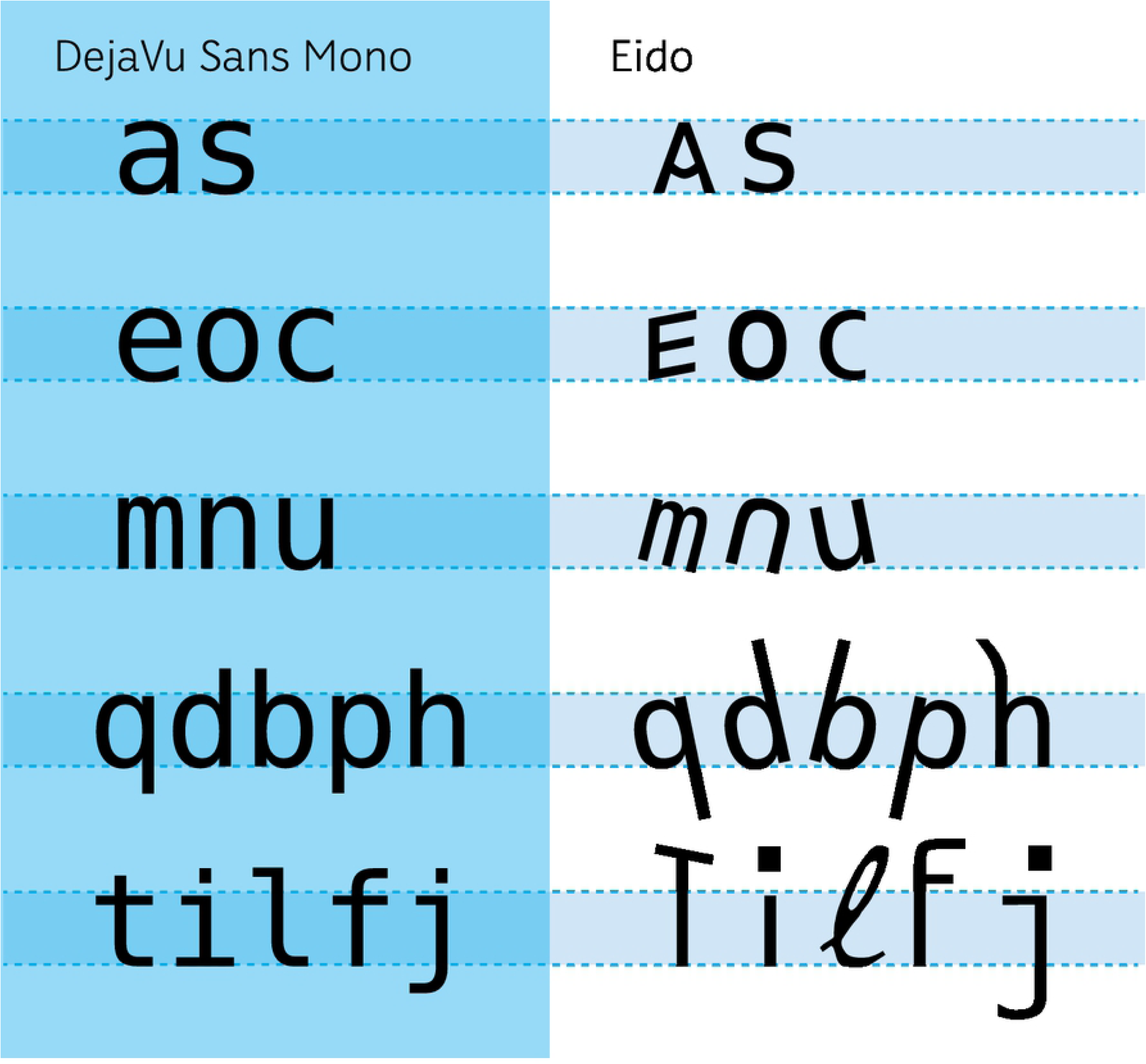
The fonts DejaVu Sans Mono and Eido as tested by Bernard et al (7). Eido is based on DejaVu and contains letter groups of mixed upper- and lowercase letters; slant to left and right; and longer ascenders and descenders.

The absence of regularity in the Eido font (Fig 2), makes it a very unusual font as a whole when letters are put together to make words, even if readers are familiar with each of the individual letter templates. Eido letters look as if they belong to different fonts mixed together, as typographic nonsense. This is also important, since previous research has showed that although readers may improve their performance by reading fonts with uncommon letter shapes, they do not like to do so (10). It also suggests that without prior practice and familiarization with the font style, the lack of font tuning would have a negative effect on reading text set in Eido. In multiple cases, font tuning has been demonstrated in central vision (11-13). This phenomenon occurs when readers recognize a sequence of letters presented in the same font faster than when presented with a mix of fonts. The effect has been shown in search tasks when readers recognize a target letter among letters of the same font compared to a mix of font styles (13), or when lexical decision is positively affected by successively presented words being set in the same font compared to switching between fonts (Cooper Black and Palatino Italic) (14) and switching between fonts of the same font family (Regular and Bold) (15). The results are interpreted as an indication that the perceptual system processes text representation by identifying the specific structures of a font and then tunes into these features (16).

This notion of feature tuning has parallels with findings on words set in miXeD cAsE. The negative effect on recognition of mixed-cased words has been shown in multiple experiments investigating lexical decision (17) and sentence reading (18, 19). By employing a lexical decision paradigm with central visual presentation, recent research has demonstrated that this mixed-case effect is unrelated to the availability of lexico-sematic information and is instead due to a lack of visual familiarity (20). The findings on font tuning (mixed-fonts) and the findings on visual familiarity (mixed-case) all suggest that as a reader is presented with a word, the perceptual system relies on prior exposure to specific visual rule sets concerning how components within a word relate to each other.

In this paper we were interested in the effect of fonts of varying inter-letter regularity styles on word recognition performance. We tested the hypothesis that low inter-letter regularity can have a negative effect on peripheral word recognition performance and tested whether some specific forms of lacking inter-letter regularity are more deleterious than others.

## 2. Font design

We designed four new fonts that are all based on the traditional font DejaVu Sans. We took great care in developing versions where the letter shapes were of familiar structures. In other words, it was essential not to reinvent the alphabet in the aim for a high degree of letter differentiation. The categories were as follows: A) The Extended category has exceptionally long ascenders and descenders, the longer extenders increasing the dissimilarity between letters such as ‘n’ > ‘h’, and ‘o’ > ‘p’. While so doing, we maintain the important modular system that typographers find essential for fluent reading (2). The modular system of the Extended font results in good inter-letter regularity. B) The Slant category is made by rotating letters to the left and right, while the letters maintain their internal relation. This rotation breaks with the inter-letter regularity, as letter pairs such as ‘h’ and ‘b’ no longer have common paths when superimposed. The lack of a modular system for the Slant font results in poor inter-letter regularity. C) The MixedCase category, defined as a mix of lower- and uppercase letters, is based on findings that mixed-case text has low visual familiarity (20), as it breaks with all typographical rules concerning the repetition of shapes (in contrast to fonts of good inter-letter regularity, ‘b’ and ‘p’ share no modules). The lack of a modular system for the MixedCase font results in poor inter-letter regularity. Three of the fonts contain only one visual category, while one contains all three categories (Fig 3).

**Fig 3.**
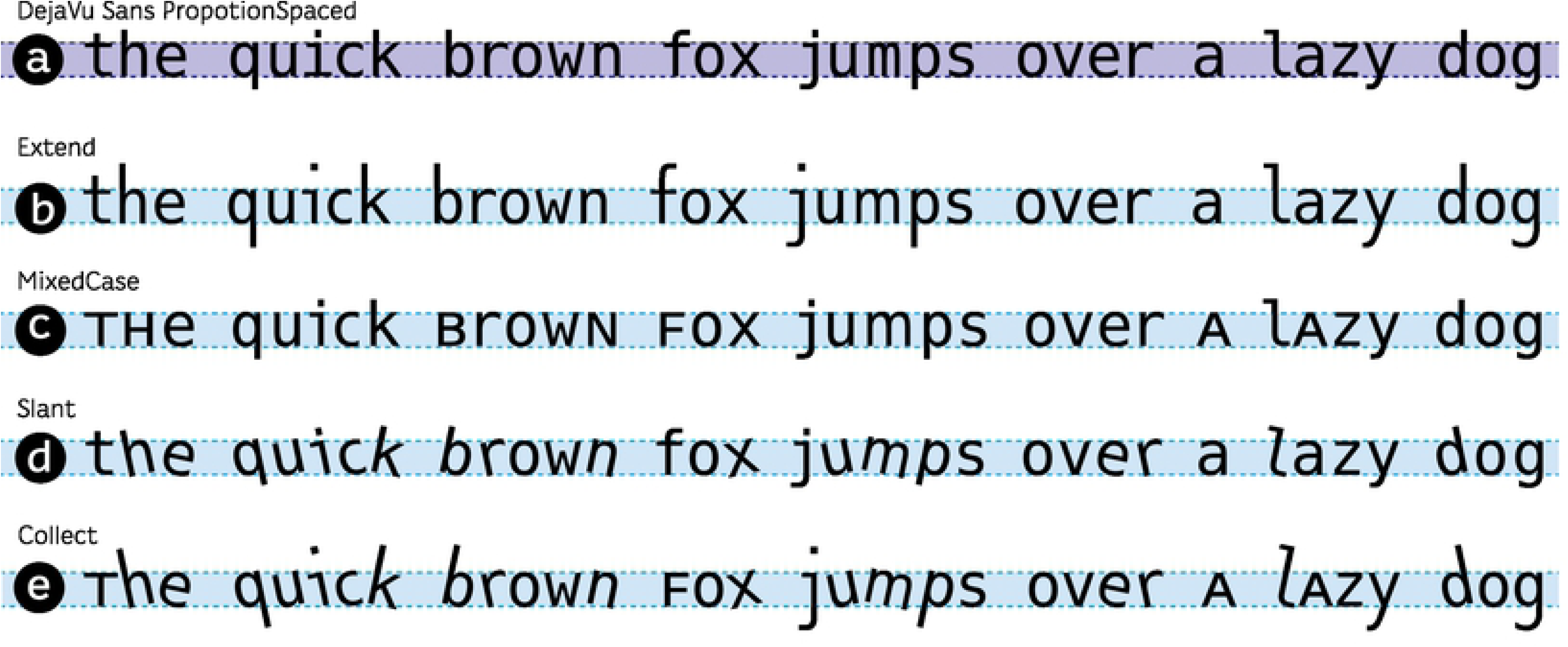
The fonts are based on the DejaVu Sans Mono font (a) and are all designed for the present investigation. The font family includes the three categories: b) Extend with exceptionally long ascenders and descender, c) MixedCase with uppercase letter shapes as x-height characters and (d) Slant with a mix of letter slant and a letter rotation of +/-12 degrees. The fourth font (e) incorporates all three categories.

All fonts were tested with the same x-height. The two fonts with long extenders (Collect and Extend) were therefore presented in a larger total vertical size than the fonts with regula-length extenders (Fig 4).

**Fig 4.**
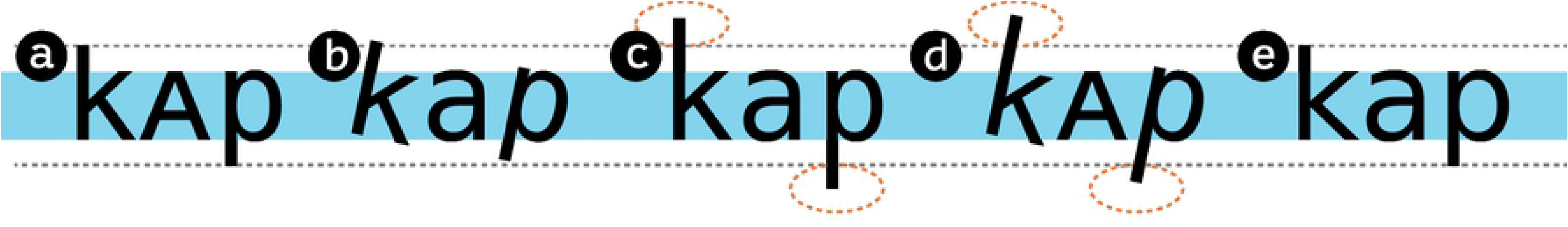
All fonts have the same x-height. Due to the long extenders, c) Extended and d) Collect take up more vertical space than the other fonts.

## 3. Experiment 1. Word recognition

We tested word recognition performance for each of the newly designed fonts and compared it to a master font in a traditional design (DejaVu Sans).

### 2.1 Subjects

The six subjects who participated in the experiments all had self-reported normal or corrected-to-normal vision. The subjects were aged 20 to 25 years (mean age 23 years), three were females, and they were recruited through the website forsoegsperson.dk. Written informed consent was obtained from the subjects after the nature of the study had been explained to them. The research complied with the Declaration of Helsinki and The Danish Code of Conduct for Research Integrity. All subjects received a gift card of DKK 300 upon completion of the experiment.

### 2.2 Apparatus

Stimuli were displayed on a 17-inch IBM/Sony CRT monitor (refresh rate = 85hz, resolution = 1024 × 768) connected to an ASUS laptop PC. Experiments were written using the software OpenSesame (21).

The experiments were carried out in a darkened room with dim lighting. Subjects were placed at a viewing distance of 50 cm from the screen. The stimuli were presented as white text on a black background.

### 2.4 Words and pseudowords

The 500 Danish words were Danish lemmas of four to six characters with a lexical frequency of 0.00002 to 0.03 percent of occurrences. The pseudowords were generated by changing one letter of existing words; care was taken so the change resulted in a pseudoword and not a new, actual word.

### 2.5 Procedure

We compared word recognition performance for the different fonts through a lexical decision task. Details of the experiment are shown in Fig 5.

**Fig 5.**
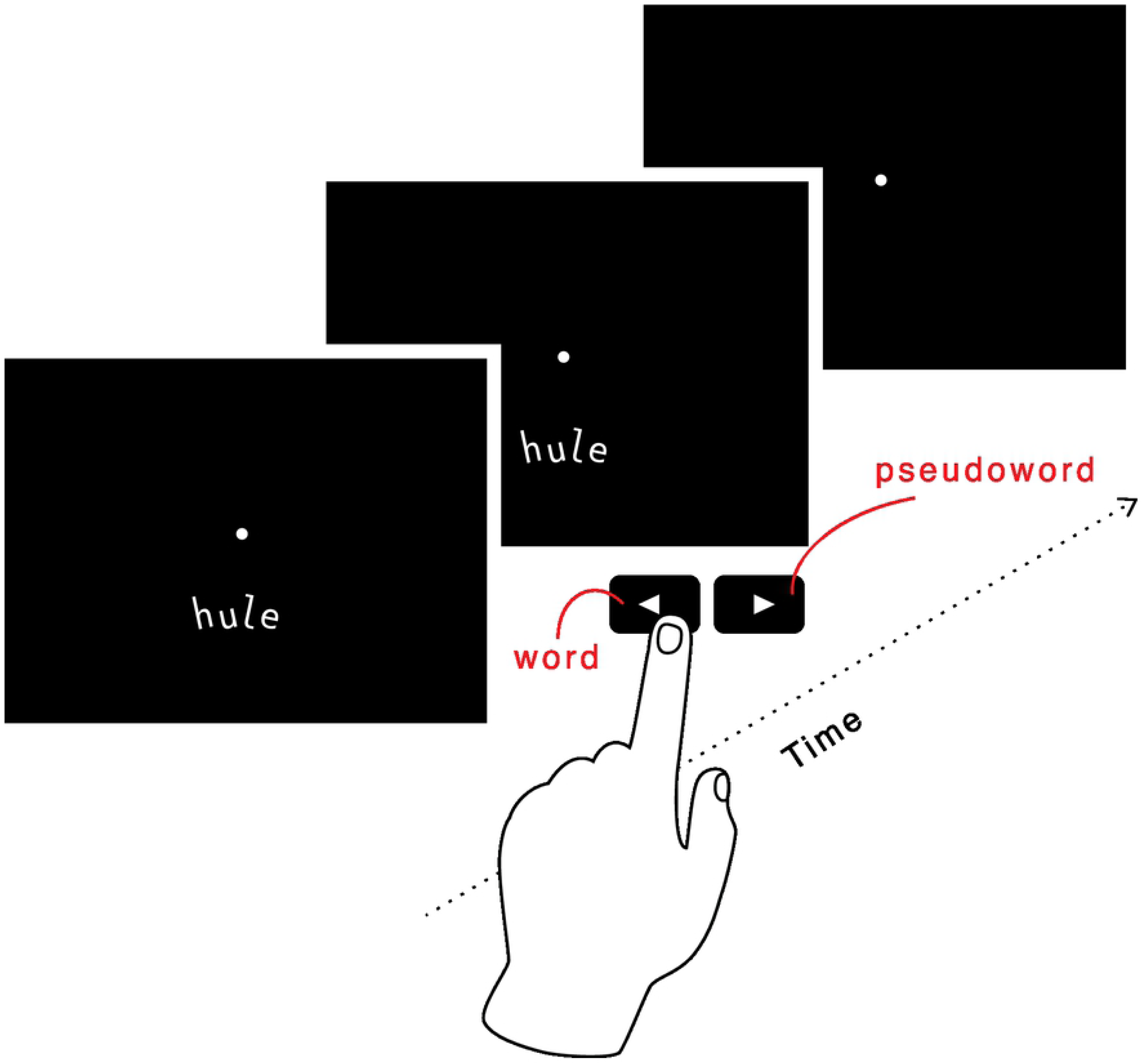
Description of the experimental protocol for the lexical decision task.

The subject was asked to fixate on a central dot while words or pseudowords were randomly presented at 10° in the lower visual field. The experimenter kept a close watch on the subject to control for steady fixation on the target dot. Trials that involved eye movements were discarded. When the subject was ready for a trial, he or she pressed the down arrow on the keyboard, after which the exposure occurred. To carry out the task, the subject had to press the left or right arrow when he or she identified a word or a pseudoword. The session lasted about two hours and consisted of nine blocks of 100 trials for each font. The blocks were presented in random order. A total of 450 words and 450 pseudowords were presented.

### 2.6 Results, Experiment 1

The results of the lexical decision task are shown in Fig 6. The DejaVu font shows the best lexical decision performance on average across subjects (lexical decision time: 0.14 ± 0.01 log ms – average ± standard error). Collect had the worst lexical decision performance on average (lexical decision time: 0.22 ± 0.02 log ms).

**Fig 6.**
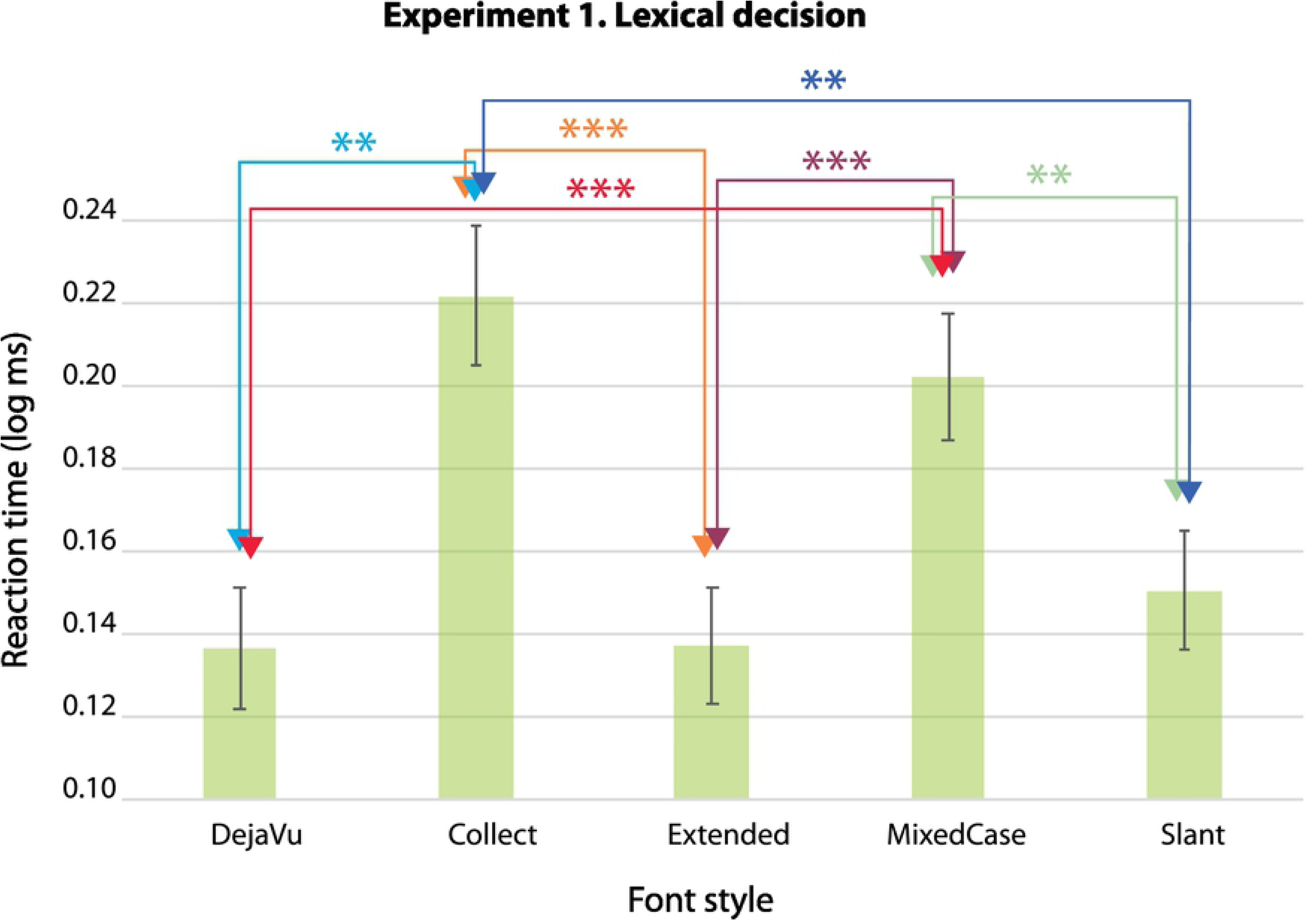
Average and standard error log response time (s) for the different fonts. P-values: ***<0.001, **<0.01, *<0.05.

The visible separation between the two groups of fonts (DejaVu, Extended and Slant vs. MixedCase and Collect is significant, as shown by a linear mixed effect with log reaction time as the dependant variable, font style as the fixed variable, and subject identity as the random variable. P-values that correspond to the differences between the different fonts are shown in Table 1.

**Table 1.** P-values for the differences between lexical task durations based on our mixed-effect model: ***<0.001, **<0.01, *<0.05

|  | DejaVu | Collect | Extended | Mixedcase | Slant |
| --- | --- | --- | --- | --- | --- |
| DejaVu |  | 0.0020** | 0.4669 | 0.0055** | 0.8654 |
| Collect | 0.0020** |  | 0.0002*** | 0.7184 | 0.0014** |
| Extended | 0.4669 | 0.0002*** |  | 0.0008*** | 0.5832 |
| MixedCase | 0.0055*** | 0.7184 | 0.0008*** |  | 0.0042** |
| Slant | 0.8654 | 0.0014** | 0.5832 | 0.0042** |  |

### 2.7 Discussion, Experiment 1

The font styles Extended and Slant both resulted in similar performances to that for DejaVu, with no statistical differences observed between the different fonts. While Extended has similar inter-letter regularity to DejaVu (both have well-functioning modular systems between letter groups, such as e-c-o, p-b-q-d and u-n-m-h), the Slant font can be considered less regular because its oblique features with multiple orientations are features that are rarely present in typical letters. The findings suggest that a poorer inter-letter regularity, which is the result from slanting letters to the left and right, does not impede word recognition performance. The same is not the case for the mixed-case fonts (MixedCase and Collect), which exhibited large and significant negative difference from the others with regard to word recognition performance. Mixed-case fonts are considered fonts with poor inter-letter regularity, as they mix two different kinds of modular systems (lower- and uppercase systems).

Based on our initial findings, we were interested in investigating letter recognition performance for the same test fonts. We wanted to ensure that our results were due to differences in inter-letter regularity, not lower-level factors, such as inter-letter confusability.

## 4. Experiment 2. Peripheral letter identification

With the same fonts and apparatus as in Experiment 1 we tested letter recognition when the stimuli were presented to subjects in trigrams (three-letter strings).

### 3.1 Subjects

Eight new subjects participated in the experiment, all self-reporting normal or corrected-to-normal vision. The subjects were aged 21 to 29 years (mean age 25 years), seven were females. As in Experiment 1, written informed consent was obtained from the subjects after the nature of the study had been explained to them. The research followed the tenets of the Declaration of Helsinki and The Danish Code of Conduct for Research Integrity. All subjects received a gift card of DKK 150 upon completion of the experiment.

### 3.2 Procedure

The task was to recognize all the letters of a trigram that was briefly presented at 10° in the lower visual field. Print size was chosen so that the recognition rate of the central letter was about 50% during a pre-trial training session of 10 trials per font. The presentation time was 200 ms. Subjects were asked to fixate on a dot centred on the screen. The experimenter kept a close watch on the eyes of the subject to control for steady fixation on the target dot.

Approximately 5% of the trials were discarded because of eye movements. The principles of the experiment are shown in Fig 7. When the subject was ready for a trial, he or she pressed the space bar on a computer keyboard, after which the stimulus exposure occurred. Following the presentation, the subject was asked to select the three stimuli letters displayed during the trial from left to right. No feedback was given to the subject. The session lasted about one hour and consisted of six blocks of 100 trials each. To avoid participants becoming familiar with the letter shapes of the fonts, each block consisted of 20 consecutive trials for each font. The font order was random for each block. For each trigram, three letters were randomly selected among the 26 letters of the alphabet.

**Fig 7.**
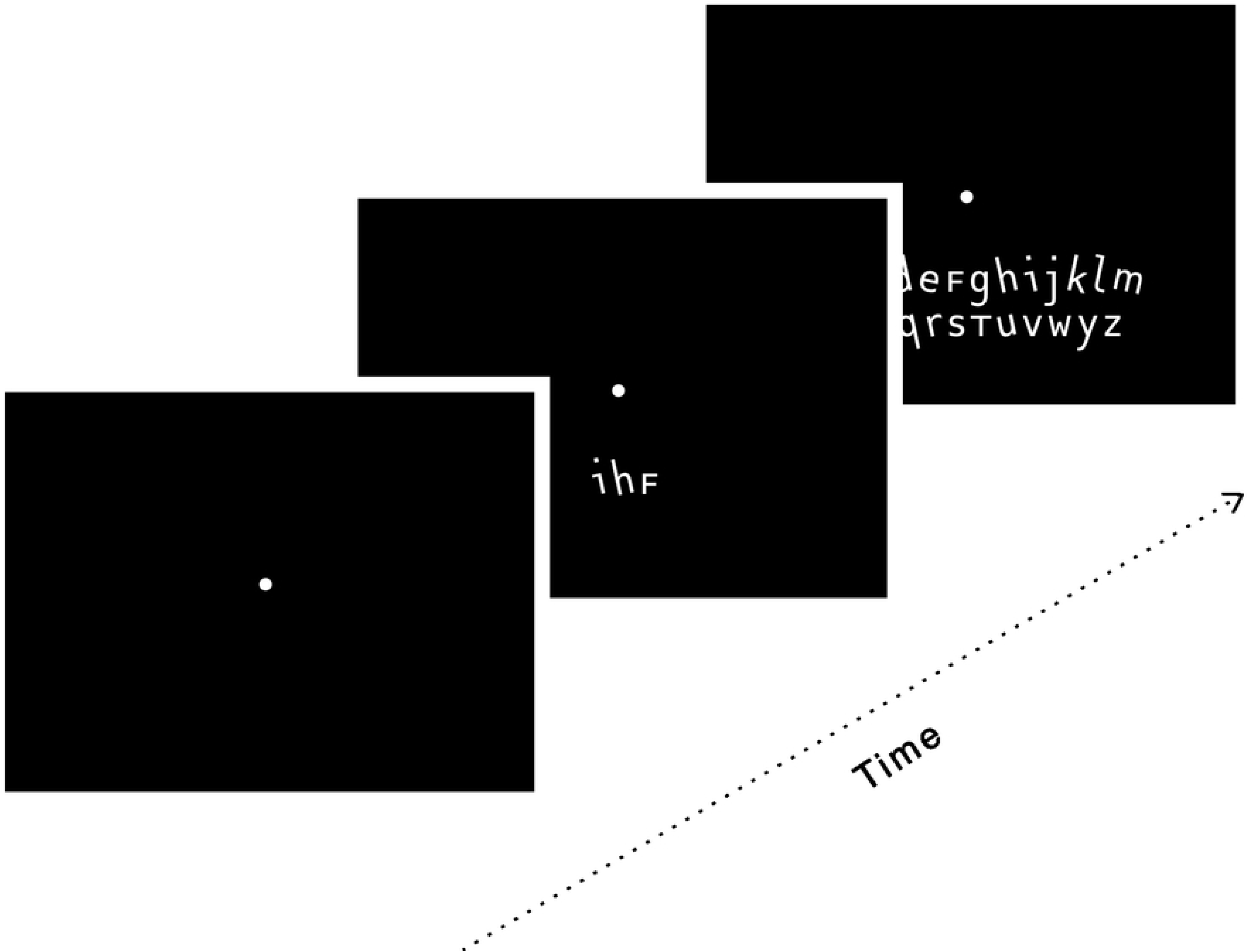
Description of the experimental protocol of trigram recognition based on a presentation time of 200 ms.

### 3.3 Results, Experiment 2

Fig 8 shows the average letter recognition rates per trigram presentation for each font. Standard errors across subjects are indicated in the figure. The traditional DejaVu font has a high recognition score (1.93 ± 0.03 letters per trigram) and on average is only inferior to the Extended font (1.97 ± 0.03 letters per trigram). By contrast, the fonts with the poorest recognition rates are the Collect, MixedCase and Slant fonts, which have an average recognition rate between 1.79 and 1.89 letters per trigram, with the Slant font resulting in the poorest performance.

**Fig 8.**
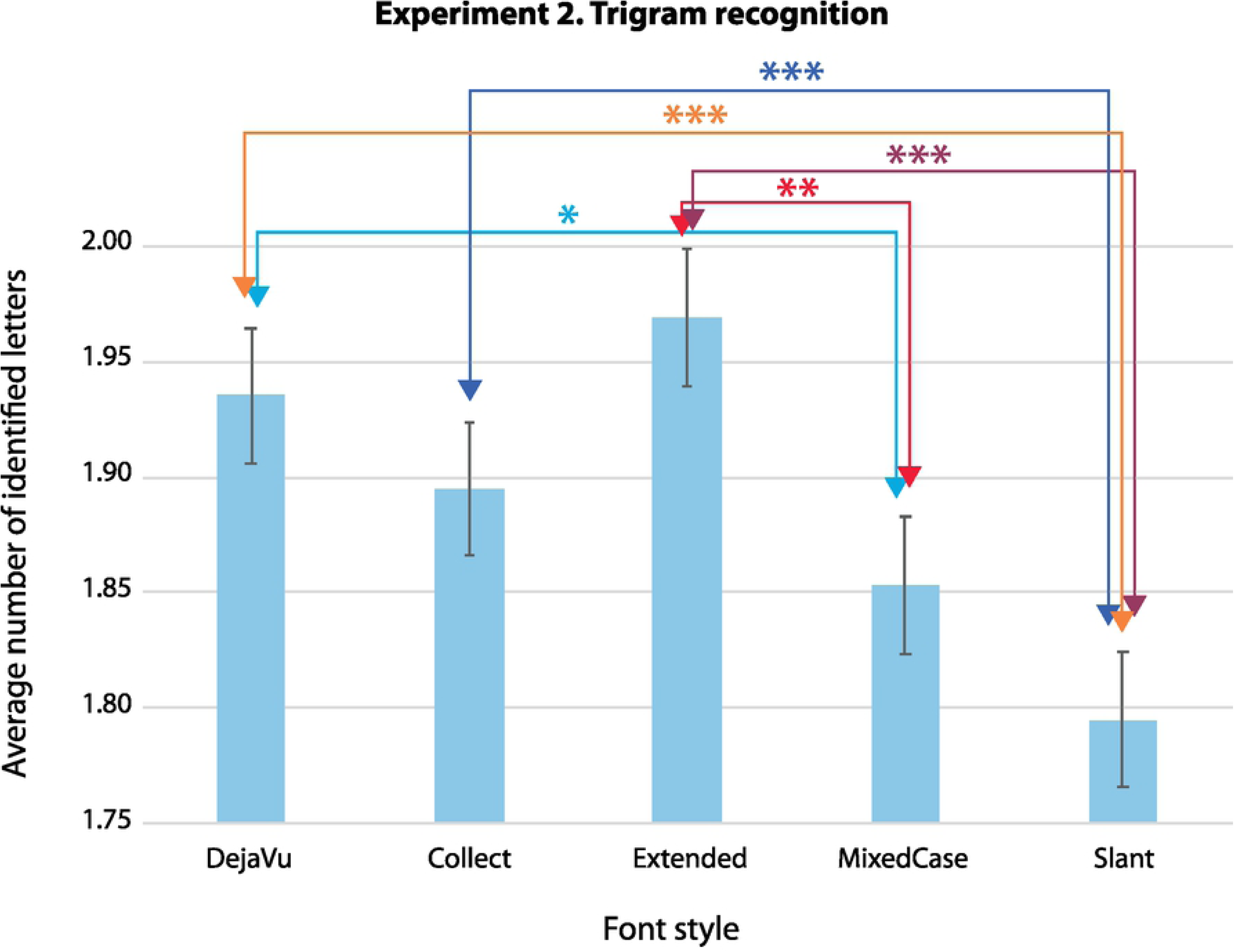
Average number of identified letters and standard error values across subjects. P-values: ***<0.001, **<0.01, *<0.05.

We ran a mixed-effect model to test whether the differences observed between the fonts were significant. The dependant variable was the number of letters correctly identified, the fixed variables were the font types, and the random variable was the subject identity. P-values that correspond to the differences between the different fonts are shown in Table 2.

**Table 2.**
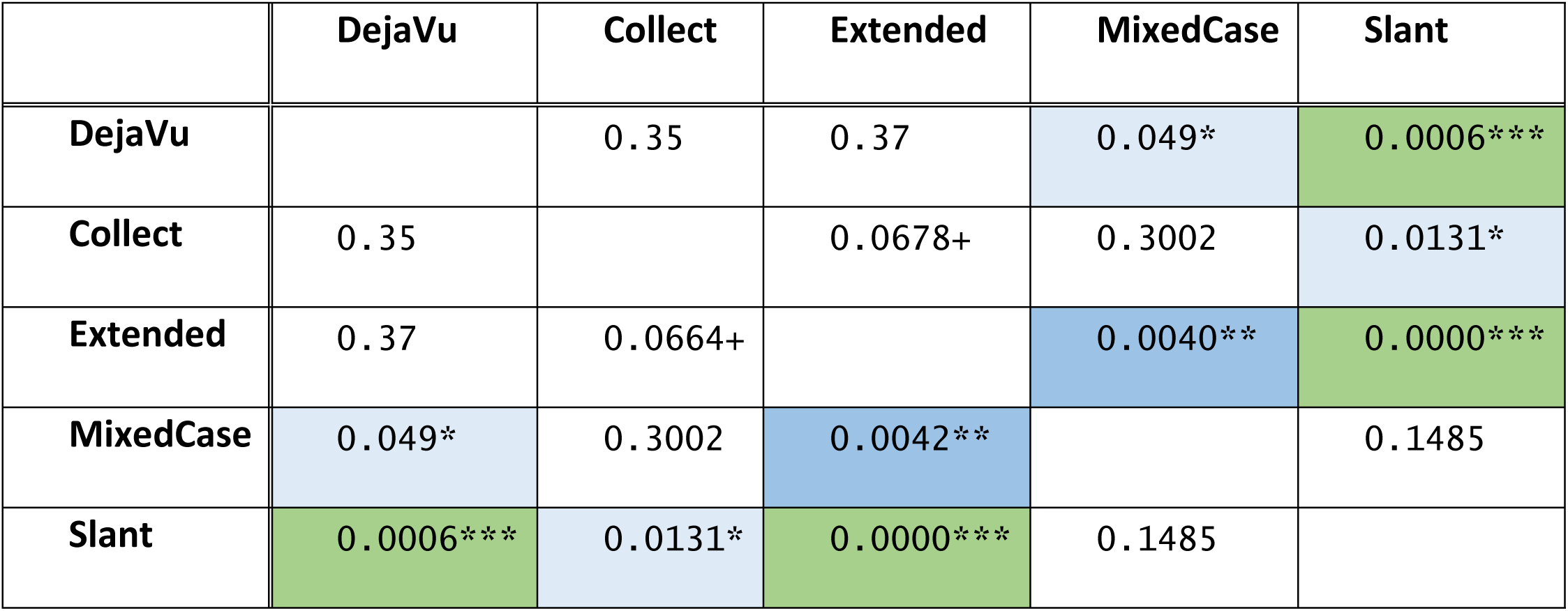
P-values corresponding to the differences between the different fonts and based on the linear mixed-effect model: ***<0.001, **<0.01, *<0.05.

|  | DejaVu | Collect | Extended | MixedCase | Slant |
| --- | --- | --- | --- | --- | --- |
| DejaVu |  | 0.35 | 0.37 | 0.049* | 0.0006*** |
| Collect | 0.35 |  | 0.0678+ | 0.3002 | 0.0131* |
| Extended | 0.37 | 0.0664+ |  | 0.0040** | 0.0000*** |
| MixedCase | 0.049* | 0.3002 | 0.0042** |  | 0.1485 |
| Slant | 0.0006*** | 0.0131* | 0.0000*** | 0.1485 |  |

The p-values confirm that the DejaVu and the Extended fonts offer a significant advantage in our peripheral letter recognition task. Statistically, Extended shows significantly better performance than the three other fonts. DejaVu is superior to two fonts, and Collect is superior to the Slant font.

Overall, the findings demonstrate that the results for letter recognition performance are very different compared to word recognition performance. More generally, the correlation between word and letter recognition performance is very poor (R2 = 0.06).

### 3.4 Discussion, Experiment 2

The Extended, DejaVu and Collect fonts had significantly higher scores with regard to letter recognition performance, meaning that they had the lowest inter-letter confusability. The Slant font had the poorest letter recognition performance followed by the MixedCase font. When we compare this with the results of Experiment 1, it suggests that what we observed in Experiment 1 (poorest performance for both fonts with mixed-case letters) cannot be due to a poor inter-letter confusability but is directly linked to the lack of regularity between the different letters.

## 4. General discussion

Our first hypothesis was that poor inter-letter regularity would impair reading performance. Our results suggest that, indeed, lack of inter-letter regularity can significantly impair peripheral word recognition performance. We showed this negative effect for two fonts (MixedCase and Collect), both mixing lowercase and uppercase letters. These fonts with the smallest inter-letter regularity (due to being a mix of lower- and uppercase modular systems) were also the fonts that significantly resulted in the poorest performances, while the fonts that had a better inter-letter regularity (DejaVu and Extended) resulted in the best performances. Interestingly, intermediary irregularity caused by tilted letters (Slant) did not significantly affect word recognition performance.

The findings of the word recognition experiment cannot be explained by letter recognition performances, as results were inconsistent between the two experiments. In the case of the Slant font, the findings show opposite results between letter and word recognition. The Slant font was the poorest-performing font with regard to letter recognition, while for word recognition it showed a similar recognition rate to the two best-performing fonts and did significantly better than the two mixed-case fonts. Our findings thus show an important limitation of the usually accepted theory that links peripheral letter and word recognition performance (22, 23). It is also possible that the lack of regularity between letters causes the disruption of word uniformity, and a consecutive decrease in word recognition performance (24).

The letters in the slant conditions were either rotated to the right or to the left or had no rotation. It appears that for letter recognition, this rotation is confusing, as it is difficult to predict the nature of the rotation for each single letter. While for word recognition, the rhythm produced by the rotations of the Slant font condition leads to greater predictability of the word components and thus makes it easier for the subjects to tune into the font structure. Our results differ from findings by Gauthier et al. (13), who compared recognition of letter trigrams where the letters were slanted to one side to the recognition of trigrams where the letters were mixed between slants to the left and right (similar to our Slant font) and found no difference in performance between the two font conditions. Since our experiment did not compare the Slant font with a font condition that only had a slant to one side, this may be the cause of the different results.

The fact that the mixed-case fonts (Collect and MixedCase) are the poorest-performing in the word recognition experiment confirms previous studies of the mixed-cased effect on foveal recognition (17-20). In the present study, we extend the findings to include peripheral vision.

In our experiment on letter recognition, only one out of the two fonts with mixed-case features was significantly outperformed by DejaVu, which indicates that the negative influence of mixed-case fonts on letter recognition is less pronounced than the impact on to word recognition. If letters within a word become too uncommon in relation to each other, subjects may have to adopt a reading strategy based on serial processing of each single letter, which is much less efficient than parallel processing drawing on orthographic lexical information (25, 26).

For both letter and word recognition, the long extenders hold an advantage (Extended). In reading situations involving smaller visual angles, a large x-height (meaning shorter extenders) is known to facilitate reading (27). However, it is possible that if the x-height is kept constant, longer extenders could also benefit reading at small visual angles. Our findings suggest that for reading situations involving peripheral reading, long ascenders and descenders may be an advantage. This is interesting, since, to our knowledge, this simple change in fonts had never been directly tested, although it seems to be an easy way to modify a font and improve letter recognition performance.

Studies into letter recognition suggest that letters are recognized by their features (6, 28-30). Viewing our findings in this perspective, the data on letter recognition suggests that as long as the letter features are identifiable, the level of inter-letter regularity is of less importance. In contrast to this, the data on word recognition suggests that word processing benefits from regularity. It is generally believed that for successful word processing, it is highly essential to be able to recognize the letters and their features (26, 31, 32); our findings add to this by demonstrating that in addition to great inter-letter dissimilarity (7), inter-letter regularity within a word also contributes to successful word recognition.

## Conclusion

We found evidence that a new factor, which we have labelled regularity, has a direct effect on word recognition performance, as fonts of great inter-letter regularity outperformed fonts of low inter-letter regularity in a peripheral word recognition task. The effect varied between letter and word recognition, so that rotated familiar letter shapes had a more negative effect on letter recognition than on word recognition, and mixing upper- and lowercase letters – which was generally detrimental – had a more negative effect on word recognition than on letter recognition. Our key finding is that between letter and word recognition, great inter-letter regularity has the most positive effect on word recognition and less on letter recognition, which shows that supplementary features can improve letter recognition, while they have a negative effect on word recognition. Our findings demonstrate that the typographic approach of working with inter-letter regularity is an important factor that needs to be considered in the design of fonts for word processing in peripheral vision.

## Acknowledgments

We are grateful to the Society for Danish Language and Literature for creating a list of orthographic neighborhood sizes for Danish words and a list of the most common lemmas, which is based on the all the entries in the Danish Dictionary. This work was supported by the Danish Council for Independent Research [grant number DFF – 7013-00039].

## References

1. Carter M. An Exercise in Versatility. In: Cabarga L, editor. Logo, Font & Lettering Bible: A comprehensive guide to the design, construction and usage of alphabets, letters and symbols: Davis & Charles; 2004. p. 200.

2. Beier S. Type Tricks: Your Persomal Guide to Typedesign: BIS Publishers; 2017.

3. Gates D. Lettering for Reproduction. New York: Watson-Guptill Publications; 1969.

4. Blokland FE, der Kunsten A. On the origin of patterning in movable Latin type: Renaissance standardisation, systematisation, and unitisation of textura and roman type 2016.

5. Beier S. Reading Letters: designing for legibility: BIS Publishers; 2012.

6. Fiset D, Blais C, Ethier-Majcher C, Arguin M, Bub D, Gosselin F. Features for identification of uppercase and lowercase letters. Psychological science. 2008;19(11):1161–8.

7. Bernard J-B, Aguilar C, Castet E. A New Font, Specifically Designed for Peripheral Vision, Improves Peripheral Letter and Word Recognition, but Not Eye-Mediated Reading Performance. PloS one. 2016;11(4):e0152506.

8. Xiong Y-Z, Lorsung EA, Mansfield JS, Bigelow C, Legge GE. Fonts designed for macular degeneration: Impact on reading. Investigative ophthalmology & visual science. 2018;59(10):4182–9.

9. Beier S, Larson K. Design improvements for frequently misrecognized letters. Information Design Journal. 2010;18(2):118–37.

10. Beier S, Larson K. How does typeface familiarity affect reading performance and reader preference? Information Design Journal. 2013;20(1):16–31.

11. Sanocki T. Visual knowledge underlying letter perception: Font-specific, schematic tuning. Journal of Experimental Psychology: Human Perception and Performance. 1987;13(2):267.

12. Sanocki T. Font regularity constraints on the process of letter recognition. Journal of Experimental Psychology: Human Perception and Performance. 1988;14(3):472.

13. Gauthier I, Wong AC, Hayward WG, Cheung OS. Font tuning associated with expertise in letter perception. Perception. 2006;35(4):541–59.

14. Walker P. Font tuning: A review and new experimental evidence. Visual Cognition. 2008;16(8):1022–58.

15. Dyson MC, Beier S. Investigating typographic differentiation: Italics are more subtle than bold for emphasis. Information Design Journal. 2016;22(1):3–18.

16. Sanocki T, Dyson MC. Letter processing and font information during reading: Beyond distinctiveness, where vision meets design. Attention, Perception, & Psychophysics. 2012;74(1):132–45.

17. Allen PA, Wallace B, Weber TA. Influence of case type, word frequency, and exposure duration on visual word recognition. Journal of Experimental Psychology: Human Perception and Performance. 1995;21(4):914.

18. Reingold EM, Rayner K. Examining the word identification stages hypothesized by the EZ Reader model. Psychological science. 2006;17(9):742–6.

19. Reingold EM, Yang J, Rayner K. The time course of word frequency and case alternation effects on fixation times in reading: Evidence for lexical control of eye movements. Journal of Experimental Psychology: Human Perception and Performance. 2010;36(6):1677.

20. Perea M, Fernández-López M, Marcet A. Does CaSe-MiXinG disrupt the access to lexico-semantic information? Psychological research. 2018:1–9.

21. Mathôt S, Schreij D, Theeuwes J. OpenSesame: An open-source, graphical experiment builder for the social sciences. Behavior research methods. 2012;44(2):314–24.

22. Bernard J-B, Castet E. The optimal use of non-optimal letter information in foveal and parafoveal word recognition. Vision research. 2019;155:44–61.

23. Legge GE, Cheung S-H, Yu D, Chung ST, Lee H-W, Owens DP. The case for the visual span as a sensory bottleneck in reading. Journal of vision. 2007;7(2):9-.

24. Rummens K, Sayim B. Disrupting uniformity: Feature contrasts that reduce crowding interfere with peripheral word recognition. Vision research. 2019;161:25–35.

25. Houpt JW, Townsend JT, Donkin C. A new perspective on visual word processing efficiency. Acta psychologica. 2014;145:118–27.

26. Coltheart M, Rastle K, Perry C, Langdon R, Ziegler J. DRC: A dual route cascaded model of visual word recognition and reading aloud. Psychological review. 2001;108(1):204–56.

27. McCarthy MS, Mothersbaugh DL. Effects of typographic factors in advertising-based persuasion: A general model and initial empirical tests. Psychology & Marketing. 2002;19(7-8):663–91.

28. Petit J-P, Grainger J. Masked partial priming of letter perception. Visual Cognition. 2002;9(3):337–53.

29. Rosa E, Perea M, Enneson P. The role of letter features in visual-word recognition: Evidence from a delayed segment technique. Acta psychologica. 2016;169:133–42.

30. Lanthier SN, Risko EF, Stolz JA, Besner D. Not all visual features are created equal: Early processing in letter and word recognition. Psychonomic bulletin & review. 2009;16(1):67–73.

31. Perry C, Ziegler JC, Zorzi M. When silent letters say more than a thousand words: An implementation and evaluation of CDP++ in French. Journal of Memory and Language. 2014;72:98–115.

32. Pelli DG, Farell B, Moore DC. The remarkable inefficiency of word recognition. Nature. 2003;423(6941):752–6.

